# High co-occurrence of invasive wetland plants and species at risk in Canada’s biodiverse Carolinian

**DOI:** 10.1101/2025.11.09.687476

**Authors:** Autumn D. Watkinson, Cailyn Carscadden, Rebecca C. Rooney

**Affiliations:** Trent School of the Environment, Trent University, 1600 West Bank Drive, Peterborough, ON K9L 0G2, Canada; Department of Biology, University of Waterloo, 200 University Avenue West, Waterloo, ON, N1L 3G1, Canada

**Author notes:** **Author Contributions using CRediT taxonomy:** ADW and RCR wrote and edited the manuscript; AWD and CC conducted data curation and visualization; RCR provided funding, project administration, and supervision. **Submitted to:** FACETS https://www.facetsjournal.com/for-authors/instructions-to-authors.

**Keywords:** endangered species, coastal wetlands, riparian habitat, dunes, beaches, threatened species, species of concern, conservation status, conservation, biodiversity, coastal marsh

## Abstract

Invasive species are a major driver of biodiversity loss, with invasive plants increasingly threatening wetland ecosystems. Southern Ontario’s Carolinian Zone, a biodiversity hotspot supporting 79% of Ontario’s non-fish Species at Risk (SAR), is especially vulnerable. Over half (54%) of these SAR rely on wetland or semiaquatic habitats, emphasizing the importance of wetland protection for their recovery. These habitats are fragmented and highly susceptible to plant invasions. To guide conservation, we conducted a spatial co-occurrence analysis of non-fish SAR dependent on wetlands and invasive wetland plant species. We identified 33 invasive species posing current (n = 26) or imminent (n = 7) threats in the Carolinian Zone. Overlap between SAR and invasive plants was greatest in Lake Erie’s coastal marshes and shallow waters, where invasions are well documented, and also in urban areas such as Toronto, Windsor, London, and Niagara, where SAR richness was unexpectedly high. Co-occurrence of SAR and invasive plants in these regions indicates that managing invasive plants in urban wetlands could directly support SAR recovery. Marsh-nesting birds, reptiles, and wetland plants were most exposed and vulnerable to habitat alteration and resource competition. Spatial analyses help pinpoint where invasive plants most threaten SAR, enabling targeted, effective management.

## Introduction

World-wide, natural ecosystems and the species they support are experiencing severe declines. More than 43,000 species (28% of species assessed) are considered at risk by the IUCN (International Union for Conservation of Nature; IUCN, 2024) and the extent of natural ecosystems has declined by 47%, on average, compared to their pre-settlement states (IPBES, 2019). These trends are echoed in Canada where about 1 in 5 species that have been assessed have some level of conservation risk (Canadian Endangered Species Conservation Council, 2022). Declines have occurred steadily over the last 150 years, and are most acute in the southern regions of Canada which have the longest history of colonization, agricultural, urban and industrial land uses, and where human populations are most dense (Kerr & Cihlar, 2004).

Invasive species, defined as those non-indigenous species that can establish and spread to cause ecological harm in geographic locations outside their historic range boundaries (Richardson et al., 2000), are one of the top threats to biodiversity (IPBES, 2023). Across Canada, invasive non-native and/or problematic native species are listed as a threat to species-at-risk in 55% of all finalized recovery strategies (McCune et al., 2013). Invasive species can threaten species-at-risk through increased competition for resources, predation, hybridization, infection, and habitat modification (Venter et al., 2006). Understanding the status of invasive, non-indigenous species and identifying areas where species-at-risk and invasive species significantly overlap could improve prioritization of conservation management actions, including monitoring for early detection and implementation of prevention and control strategies.

The Carolinian Zone of Ontario, Canada is a biodiversity hotspot that has been significantly impacted by habitat loss and degradation. This region covers about 22,000 km^2^in the most southwestern region of Ontario; about 2% of Ontario’s land cover. Despite its relatively small area, the Carolinian contains about 25% of Canada’s human population (Kraus & Hebb, 2020). Fully 86% of pre-settlement natural habitat has been lost and remnant natural areas within this region face extreme pressure due to anthropogenic activity. Remnant habitat is generally found in small, highly fragmented and degraded states (Kraus & Hebb, 2020). Despite this, the Carolinian has an extremely high species richness: supporting 2,200 species of herbaceous plants and 70 species of trees (Ontario Ministry of Natural Resources, 2000). Close to 400 species of birds have been recorded, and many faunal species are found nowhere else in Canada (Ontario Ministry of Natural Resources, 2000). Many of these species rely on wetland habitat to fulfill at least one life history requirement (e.g., foraging, nesting, rearing offspring). Wetlands are one of the most imperiled ecosystems in the Carolinian. In southwestern Ontario, including the Carolinian zone, over 85% of the original wetlands have been drained or filled in (Ducks Unlimited, 2010).

The high human population density and the connectivity of many Carolinian wetlands to the Great Lakes puts them at higher risk of biological invasion by aquatic species that can spread through natural dispersal, hitchhiking/fouling, and intentional release (Davidson et al., 2021). Invasive plants are a particular threat to flora and fauna in wetlands, riparian, and other semi-aquatic habitat, where they may alter habitat conditions dramatically through a variety of mechanisms that affect community composition and ecosystem properties, including productivity, nutrient cycling, and hydrology (Peller and Altermatt, 2024; Zedler and Kercher, 2004). Invasive plants may compete directly with native plants through allelopathy (Hierro & Callaway, 2003) and smothering or strangling (Gould & Gorchov, 2000), or indirectly by monopolizing aboveground (Funk, 2013) or belowground resources (Broadbent et al., 2018) facilitated by rapid growth or a high reproductive potential (Blossey & Notzold, 1995). They may also threaten native species through hybridization (Gaskin, 2017), increased pest or disease transmission (Denóbile et al., 2023), or by modifying the environment, making it less suitable for native plants (e.g., altering hydrology or nutrient regimes; Devine & Fei, 2010). Similarly, invasive plants may reduce the fitness of native faunal species by altering their habitat. For example, by increasing erosion (Hardwick et al., 2025), deoxygenating the water (Shillinglaw, 1981), altering food availability (Villamagna & Murphy, 2010), or simply by creating impenetrable stands or mats of vegetation (Warren et al., 2001).

Our objective was to better understand the spatial distribution and overlap of species at risk and invasive wetland plants in the Carolinian zone of Canada. We sought to: 1) quantify and map the distribution of invertebrate and non-fish vertebrate species at risk that depend on semi-aquatic or wetland habitat; 2) quantify and map the distribution of relevant invasive plant species capable of occupying semi-aquatic or wetland habitat; and 3) assess the ecological impact and threat to species at risk for each invasive plant species identified. From this analysis, we draw insights about the dominant mechanisms by which invasive plants threaten wetland dependent species at risk and identify regions of the greatest threat and vulnerability to inform biodiversity conservation decisions.

## Methods

### Study Area and Ecosystems

The Carolinian Zone covers approximately 22,000 km^2^ of Ontario’s far southwest (Figure 1) and is in the Great Lakes Watershed. The Carolinian is bounded by Lake St Clair and Lake Huron to the west, Lake Erie to the south, and Lake Ontario to the east. Several large rivers are found within the Carolinian, including the Grand, Thames, Detroit, and Humber Rivers (all designated Canadian Heritage Rivers), Credit, Niagara, and Sydenham Rivers, and Big Creek, although all their floodplains are heavily modified. Only 1% of this region is within conserved or protected areas (national parks, provincial parks, protected areas) (Krauss & Hebb, 2020). The area is largely under private land ownership and has a high human influence index compared to the rest of Canada (Sanderson et al., 2002). The climate of the Carolinian zone is classified as Humid High Moderate Temperate (Ecoregions Working Group, 1989). Mean annual temperature is between 6.3 and 9.4°C, with mean annual precipitation of 776-1,018 mm. Mean summer rainfall is 196-257 mm (Ecoregions Working Group, 1989).

**Figure 1.**
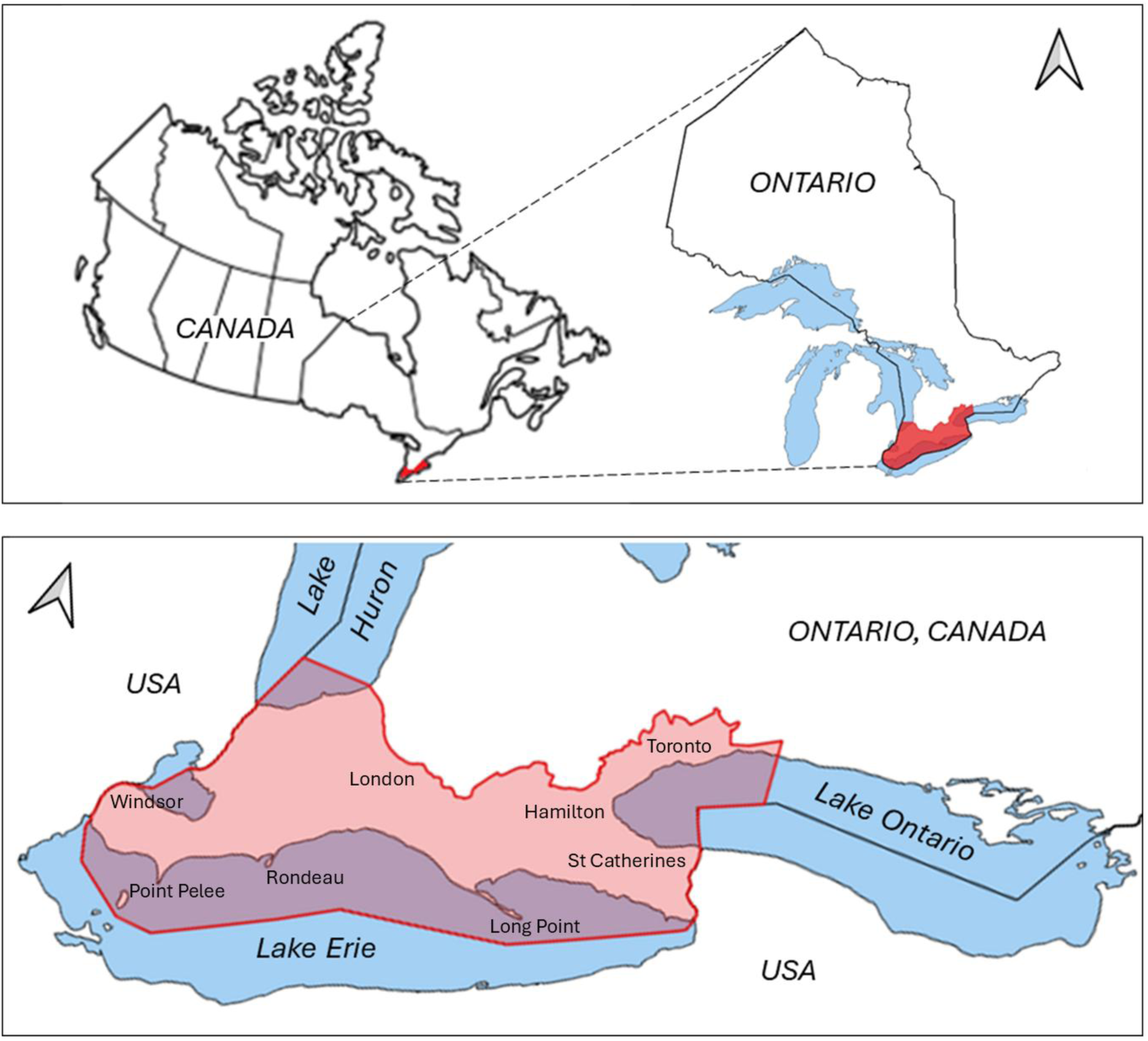
The Carolinian zone shown (red) in southern Ontario and is bordered Lake Erie, Lake St. Clair, Lake Huron, and Lake Ontario. Major urban centres and conservation lands are labelled.

We focused on wetlands (marshes, bogs, fens, swamps, shallow open water) and semi-aquatic habitats, including riparian areas, beaches, and coastal dunes that fringe open water. Wetlands were defined following the Ontario Wetland Evaluation System (Government of Ontario, 2022) as “lands that are seasonally or permanently flooded by shallow water as well as lands where the water table is close to the surface; in either case the presence of abundant water has caused the formation of hydric soils and has favoured the dominance of either hydrophytic or water tolerant plants.” Although most wetlands that existed in the region have been destroyed, some large, intact coastal marshes still exist, including the Lake Erie coastal marshes (e.g., Point Pelee, Rondeau Bay, Long Point, and Turkey Point), where the largest diversity of flora and fauna in the Carolinian region exists (Ball et al., 2003). Beaches and coastal dunes are typically sand-dominated areas found between the low-water line and upland. Though beaches and coastal dunes are not typically counted as wetland habitat, they are vulnerable to invasion by non-native wetland plants, exposing beach- or dune-dependent species at risk to threats from invasive wetland plants. For example, one of the greatest threats to the Fowler’s toad (*Anaxyrus fowleri*) is the invasive wetland grass *Phragmites australis* taking over its beach habitat and coastal dune hibernation grounds (Greenberg and Green, 2013). Riparian areas along creeks, streams, and rivers were also included, which provide important habitat connectivity for native flora and fauna, but also can create corridors facilitating the spread of invasive plants (Gregory et al., 1991). Semi-aquatic habitats constructed for purposes other than wetland conservation (e.g., stormwater management ponds, sewage lagoons, water treatment ponds) were excluded.

### Identification and Quantification of Species at Risk

We developed our list of species at risk through a series of screening steps. First, we combined all listed species at risk in Ontario (SARO) with the species listed on Schedule 1 of the federal Species At Risk Act (SARA; S.C. 2002, c. 29) that occur in Ontario. Species at Risk in Ontario were retrieved from the Species At Risk In Ontario List (O. Reg. 230/08) in November 2024. Species listed on Schedule 1 of the Species at Risk Act were retrieved from the Government of Canada species at risk public registry in November 2024 (Government of Canada, 2024). Although invasive plants can impact non-fish and fish species at risk alike, our focus in this study was on non-fish species as fish conservation in Canada is primarily governed by the Fisheries Act (Government of Canada, 2025) and falls under the jurisdiction of The Minister of Fisheries and Oceans. Coupled with economic considerations when managing commercial fish species, this leads to additional management complexities beyond the scope of our assessment.

Second, we filtered this list to retain only species that occur within the Carolinian Zone. Provincial and federal Recovery Strategies and COSEWIC (Committee on the Status of Endangered Wildlife in Canada) status reports were used to obtain distribution and habitat information for each listed species. We also used species at risk distribution data obtained from Ontario’s National Heritage Information Centre (NHIC) to ensure species were not excluded from the list based solely on distribution and habitat descriptions found in the status reports and recovery strategies.

Third, we further filtered the list to retain only species that use wetland, riparian, beach, or coastal dune habitats. We defined species at risk habitat to include breeding grounds, nursery or rearing habitat, hibernation grounds, foraging habitat, habitat critical to migration, and any other areas on which species depend directly or indirectly to carry out their life processes or any habitat where the species formerly occurred and has the potential to be reintroduced. This resulted in a list of all federally and/or provincially listed species at risk that rely on wetland, riparian, beach, or coastal dune habitat within the Carolinian zone for at least one key life process or stage, which we used in our threat analysis and mapping.

### Current and Imminent Threats from Invasive Plants

We used peer-reviewed (Davidson et al., 2021), government, and NGO lists (e.g., Great Lakes Aquatic Nonindigenous Species Information System, Ontario’s Invading Species Awareness Program, Ontario Invasive Plant Council) of invasive species and public databases (e.g., EDDMaps, iNaturalist) to identify invasive plant species that are currently found within the Carolinian. We characterized these as current threats. We also incorporated those with occurrences near the borders of the Carolinian as imminent threats. The same lists and databases were used to compile a description of each species’ growth form (i.e., terrestrial, emergent, submerged, free-floating), impact on species at risk habitat (e.g., displacement, alteration), and mechanisms of impact (e.g., resource monopolization, altered hydrology). This information was used to assess ecological impact and threat to species at risk for each invasive species identified. For all invasive plants identified, we also assigned environmental impact ratings; we used GLANSIS (Great Lakes Aquatic Nonindigenous Species Information System; GLANSIS, 2025) ratings for aquatic invasive plants and the Minnesota Invasive Species Advisory Council (2019) ratings for terrestrial invasive plants.

### Mapping the Spatial Co-occurrence of Species At Risk and Invasive Plants

#### Species At Risk Data

Occurrence data for species at risk in Ontario was obtained from the National Heritage Information Centre (NHIC) in September 2022 (Government of Ontario, n.d.), which is responsible for tracking species at risk across the province. Submitted observations are reviewed by expert biologists before being added to the database. We restricted the species at risk occurrence data we obtained from NHIC to exclude species not observed in over 20 years (circa 2002) as observations of species at risk that are more than 20 years old are often considered historical by conservation data centres, like those affiliated with NatureServe Canada (Environment and Climate Change Canada, 2024).

#### Invasive Plant Occurrence Data

A query using the EDDMaps Advanced Query Tool (EDDMapS, 2024) was conducted on January 8, 2024 to obtain occurrence and abundance data on the invasive plant species of interest. All observations entered into EDDMapS are verified by a local or state/provincial expert to ensure identifications are accurate. Query parameters included infestation status (positive, treated), country (Canada), province (Ontario), observation dates (01/01/2014– 01/01/2024), record type (current), and species. A ten-year time frame was used to reflect the dynamic nature of invasive plants and the need for recent data for effective management and control efforts.

#### Choropleth Creation

Choropleths were created using the free and open-source mapping software, QGIS (version 3.32.2). A 5 km x 5 km grid was applied to the Carolinian bounded area using the create grid function (research tools). A layer showing SAR species richness (number of species observed) and a layer showing SAR observations (number of individuals observed) for each taxonomic group (e.g., amphibians, insects, reptiles) were generated from the filtered NHIC SAR data using the count points in polygon function (analysis tools). A layer showing invasive plant observations (number of individuals observed) was generated from the EDDMaps distribution data using the count points in polygon function (analysis tools). The data in each of these layers was visualized with a graduated symbology after dividing the data into 3-5 groups of equal intervals (e.g., 9 total observations were made, data was divided into 4 classes: 0, 1-3, 4-6, 7-9). The observation (abundance) layer for invasive plants was individually overlaid by each layer created for the SAR data using the layer rendering function and the ‘multiply’ layer blending mode. A bivariate legend was generated using the bivariate legend plugin.

## Results & Discussion

### Species At Risk Occurrence & Habitat Use

Out of 250 species at risk found in Ontario (both provincially and federally listed), 29 were fish species that were excluded from our study. Of the remaining 221 species assessed in our study, 175 (79%) occur in the Carolinian. Of those, 95 (54%) use wetland habitat for some critical life stage or resource need, reflecting the strategic significance of wetland habitat protection and restoration in biodiversity conservation for Ontario. The proportion of species listed as endangered (59-60%), threatened (12-19%) and special concern (18-21%) was fairly consistent between provincial (SARO) and federal (SARA) conservation status designations (Figure 2). However, status rankings did not necessarily match for individual species, and 11 species listed provincially were not listed federally, whereas 2 species on the SARA list were not included in the SARO list. Deviations in conservation status typically reflect where the species’ population trends in Ontario differ from the rest of the country. By taxonomic group, plants had the most at-risk species (28), followed by molluscs (18), birds (17), reptiles (12), amphibians (7), insects (7), and mammals (4) (Figure 3).

**Figure 2.**
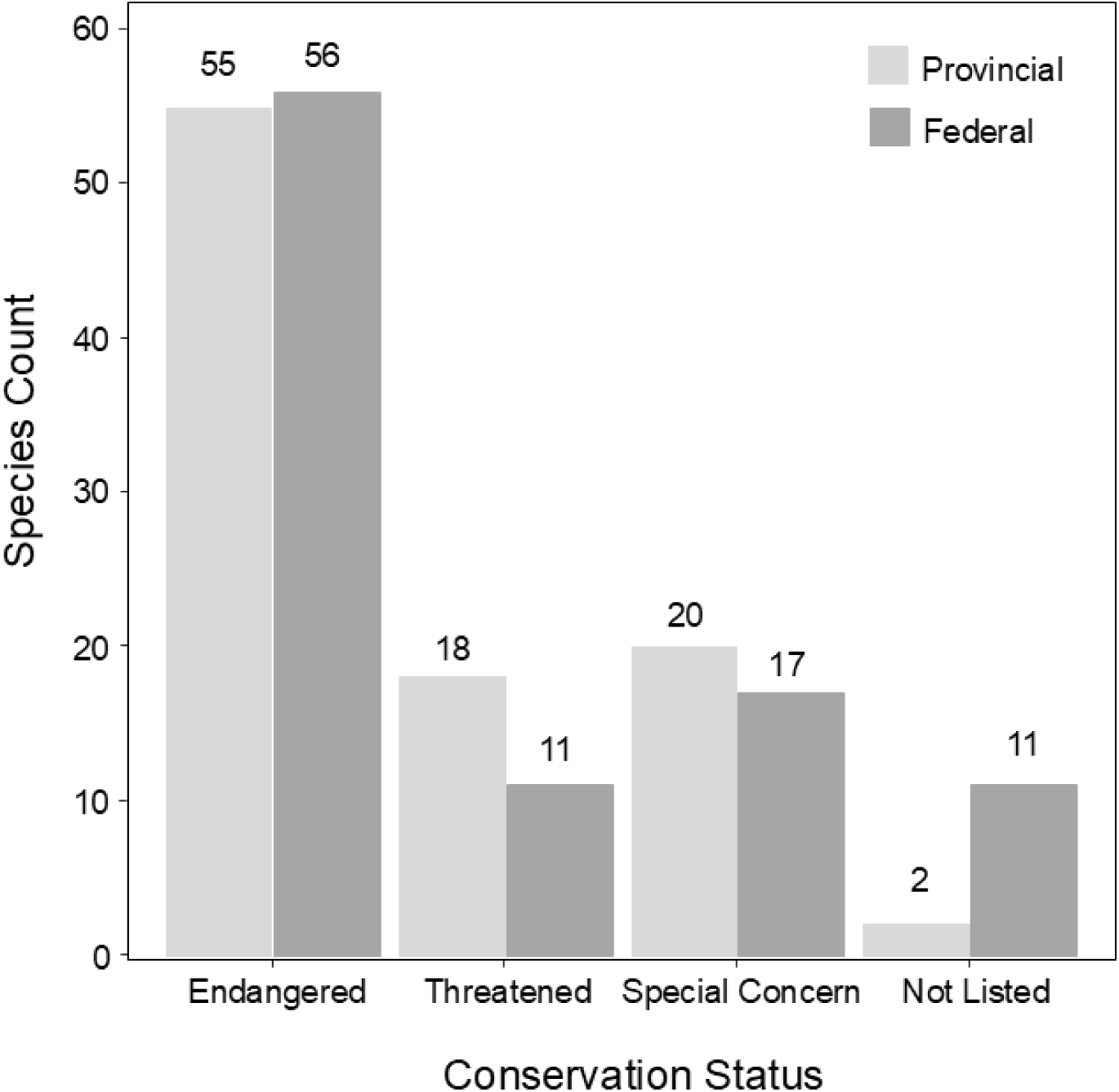
Provincial and federal conservation status of species at risk that use wetland, riparian, beach or coastal dune habitat and occur in the Carolinian, n=95.

**Figure 3.**
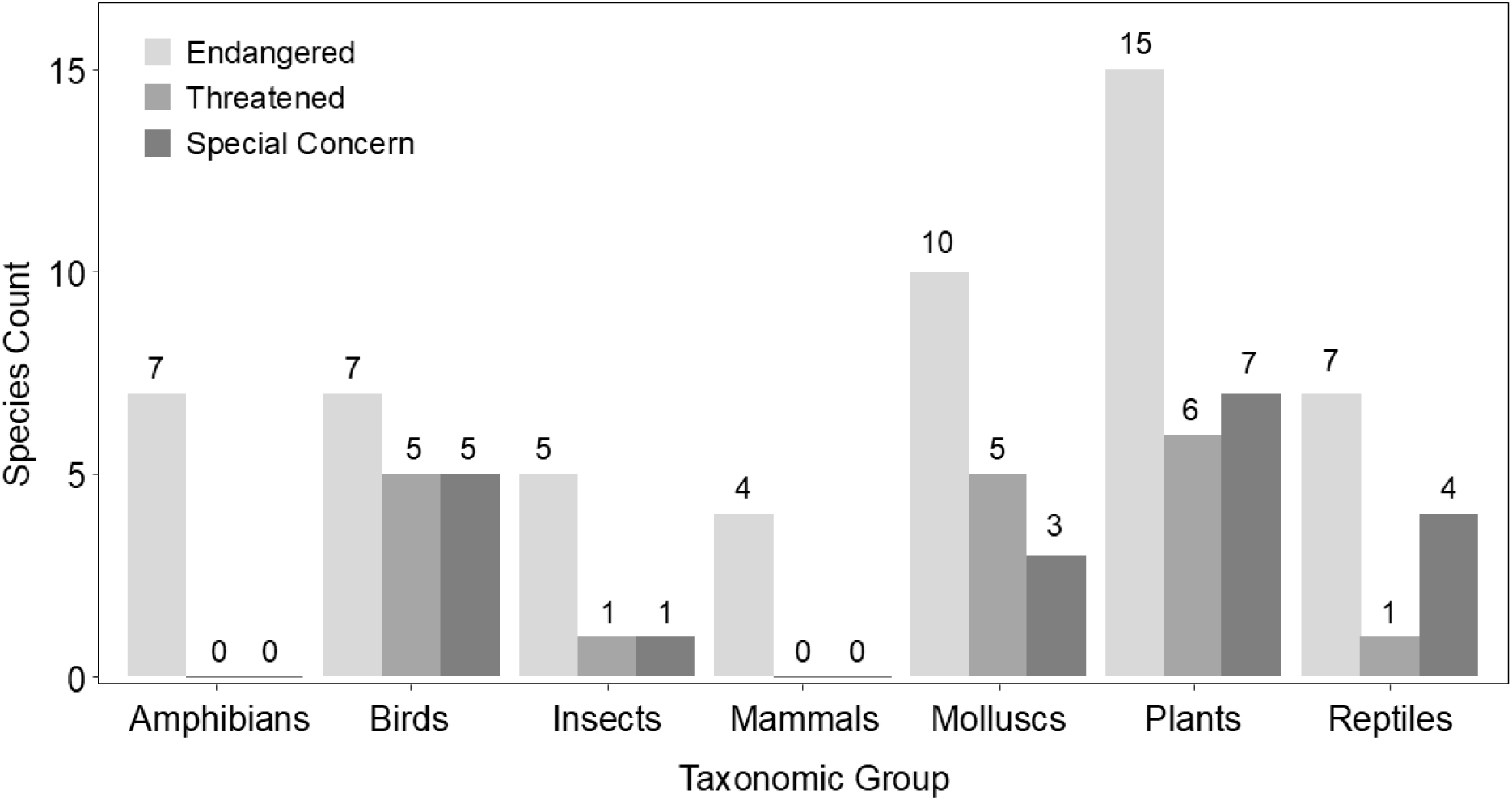
Conservation status in Ontario (SARO) for each taxonomic group of species at risk, n=95.

Many species used more than one wetland habitat type (Table 1). Unspecified wetland habitats (i.e., “wetlands”) as opposed to specific wetland classes like “bogs” or “marshes” were the most frequently described habitat used by species at risk (Figure 4). This lack of habitat specificity in many federal and provincial status reports and recovery strategies presented a challenge in interpretation as threats to species at risk from invasive plants vary substantially across wetland classes. For example, emergent invaders like *P. australis* and *Typha × glauca* often invade marshes which have shallow water habitat (Boers & Zedler, 2008; Robichaud & Rooney 2017, 2022; Tuchman et al., 2009; Zedler and Kercher, 2004), while submerged species like *M. spicatum* are more problematic in open water and slow-flowing streams (Smith and Barko, 1990, e.g., Madsen et al., 1991). Without more detailed habitat delineation, management actions risk being misdirected or inefficient. We recommend status reports and recovery strategies use standardized, ecologically meaningful habitat classifications for wetlands, like the Canadian Wetland Classification System (National Wetlands Working Group, 1997).

**Figure 4.**
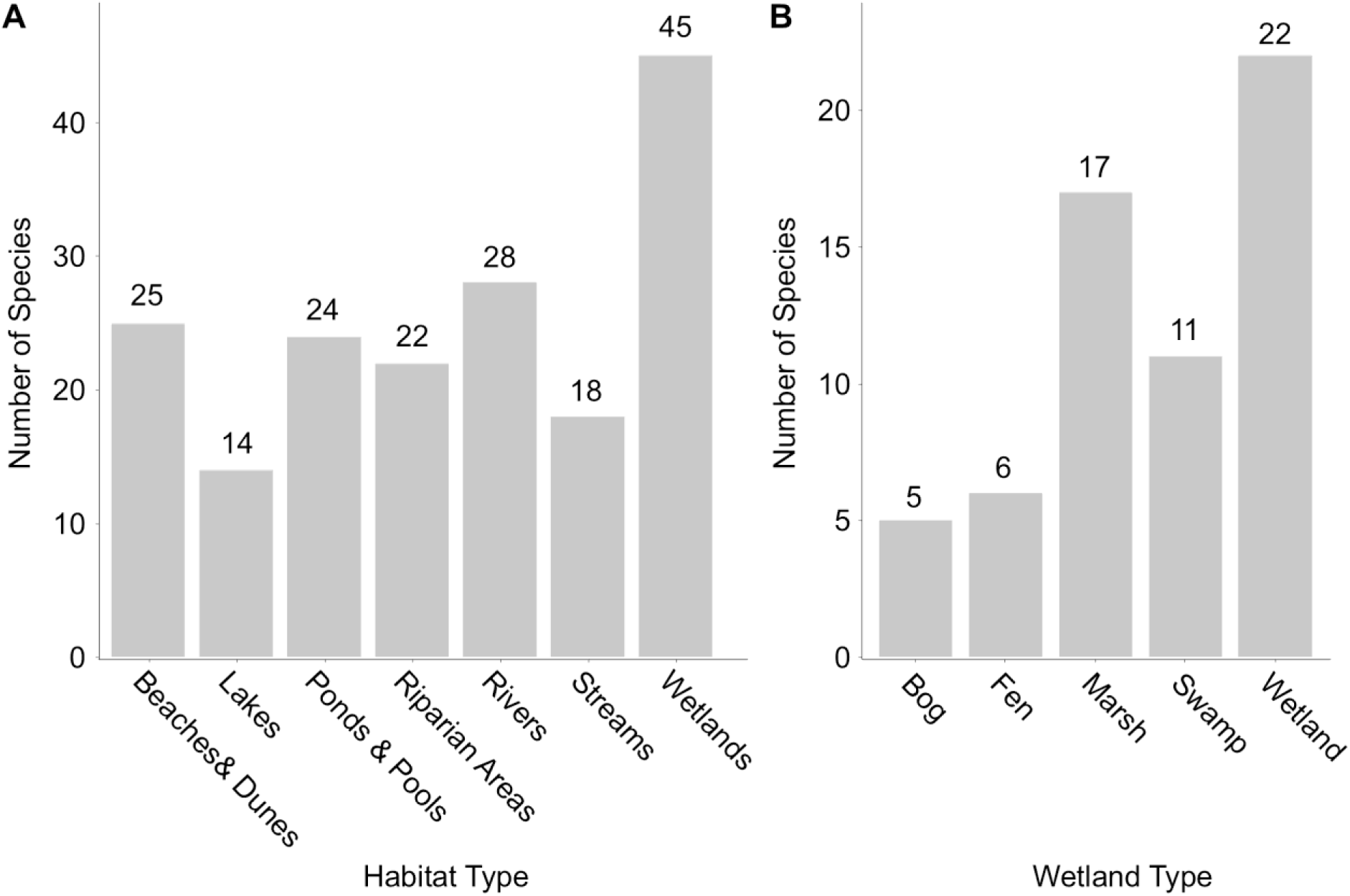
Number of species at risk using each habitat type (A). Number of species using specific wetland types (B). Many species use multiple habitats and wetland types and so the sum of column totals exceeds the 95 species at risk in our study.

**Table 1.** Federal (SARA) and provincial (SARO) conservation status (endangered (EN), threatened (TN), special concern (SC)) and use of wetlands.

| Species Name | Common Name | SARO | SARA | Habitat Description and Use |
| --- | --- | --- | --- | --- |
| Amphibians |  |  |  |  |
| *Ambystoma jeffersonianum* | Jefferson salamander | EN | EN | Upland ponds are required for breeding. |
| *Ambystoma laterale* – (2)<br>*jeffersonianum* | Jefferson salamander<br>dependent population | EN | EN | Upland ponds are required for breeding. |
| *Ambystoma laterale* – *texanum* | Small-mouthed salamander<br>dependent population | EN | EN | Upland ponds are required for breeding. |
| *Ambystoma texanum* | Small-mouthed salamander | EN | EN | Upland ponds are required for breeding. |
| *Anaxyrus fowleri* | Fowler's toad | EN | EN | Wetlands and beaches are used for all habitat requirements. Eggs and tadpoles need sparsely vegetated, still-water ponds. Juvenile and adults inhabit sandy riverside and lakeshore habitats and need sand dunes for hibernation, open beach for feeding. |
| *Desmognathus fuscus* | Northern dusky salamander | EN | EN | Always close to small groundwater-fed streams, seeps, and springs. |
| *Desmognathus ochrophaeus* | Allegheny mountain dusky<br>salamander | EN | EN | Found in or near forested small streams, springs, or seeps. |
| Birds |  |  |  |  |
| *Asio flammeus* | Short-eared owl | TN | SC | Occur in a variety of open native habitats, including bogs, marshes, wetlands, estuaries. Suitable habitat tends to be based on having a reliable source of small mammal prey. |

| Species Name | Common Name | SARO | SARA | Habitat Description and Use |
| --- | --- | --- | --- | --- |
| *Calidris canutus rufa* | Red knot rufa subspecies | EN | EN | Coastal areas with extensive sandflats, mudflats, and rocky flats are used for feeding. |
| *Chaetura pelagica* | Chimney swift | TN | TN | Aquatic and wetlands are used for feeding on flying insects. |
| *Charadrius melodus* | Piping plover | EN | EN | Dry sandy or gravelly beaches are exclusively used for nesting. Feeds on insects and small crustaceans between beach and the water's edge. |
| *Chlidonias niger* | Black tern | SC | NL | Shallow marshes are used for nesting. |
| *Chordeiles minor* | Common nighthawk | SC | SC | Wetlands and waterways are used during breeding and during migration to increase foraging efficiency on flying insects. |
| *Empidonax virescens* | Acadian flycatcher | EN | EN | Nests often found over streams, generally birds are found near ravines and swamps. |
| *Euphagus carolinus* | Rusty blackbird | SC | SC | Wetlands (flooded forests, scrub along the edges of lakes, rivers and streams, and beaver ponds) are used as winter habitat and during migration. |
| *Falco peregrinus* | Peregrine falcon | SC | NL | Tall, steep cliff ledges close to large bodies of water are used for nesting. |
| *Icteria virens* | Yellow-breasted chat | EN | EN | A wide variety of early successional habitats are used, including areas near water bodies (streams, ponds, swamps) and, occasionally, wet willow-ash-elm thickets bordering wetlands. |
| *Ixobrychus exilis* | Least bittern | TN | TN | Found in a variety of wetland habitats. Nests above the water line in stands of dense vegetation, almost always built near open water, which is needed for foraging. |
| *Melanerpes erythrocephalus* | Red-headed woodpecker | EN | EN | Found in a variety of treed habitats, including riparian forests, wetlands, and beaver ponds. |

| Species Name | Common Name | SARO | SARA | Habitat Description and Use |
| --- | --- | --- | --- | --- |
| *Parkesia motacilla* | Louisiana waterthrush | TN | TN | A riparian-obligate species, it is usually found in steep, forested ravines with fast-flowing streams or less frequently inhabits deciduous swamps having large pools of open water. It nests along stream banks. |
| *Phalaropus lobatus* | Red-necked phalarope | SC | SC | Lives in coastal and inland marshes where it feeds in shallow ponds. During migration and in the winter, it is always near water, including freshwater ponds, lakes, ditches, or lagoons. |
| *Protonotaria citrea* | Prothonotary warbler | EN | EN | Nests in the trunks of dead or dying trees standing in or near flooded woodlands or swamps. |
| *Rallus elegans* | King rail | EN | EN | Found in densely vegetated freshwater marshes with open shallow water. Sometimes found in smaller isolated marshes, but most seem to prefer larger, coastal wetlands. |
| *Riparia riparia* | Bank swallow | TN | TN | Many nests are on banks of rivers and lakes. Breeding occurs in a wide variety of low-elevation habitats, including lake bluffs and stream and riverbanks. |
| Insects |  |  |  |  |
| *Coccinella novemnotata* | Nine-spotted lady beetle | EN | EN | Lives in a wide variety of areas including riparian areas. |
| *Coccinella transversoguttata* | Transverse lady beetle | EN | SC | Lives in a wide variety of areas including riparian areas. |
| *Danaus plexippus* | Monarch | SC | EN | Caterpillars feed exclusively on milkweed plants (*Asclepias* spp.), and breeding habitat is confined to places where milkweeds grow, including wetlands. |
| *Gomphus quadricolor* | Rapids clubtail | TN | EN | Typically found in clear, cool medium-to-large rivers with gravel shallows and muddy pools. Larvae occupy quiet, muddy pools. Adult females typically inhabit forests along riverbanks and only visit shallows and pools when they are ready to mate and lay eggs. |

| Species Name | Common Name | SARO | SARA | Habitat Description and Use |
| --- | --- | --- | --- | --- |
| *Prays atomocella* | Hoptree borer | EN | EN | Dependent on its sole larval host plant, Common Hoptree, which occurs on shorelines of Lake Erie. |
| *Stylurus amnicola* | Riverine clubtail | EN | EN | Found in and near streams and rivers with sandy, muddy, or gravelly beds. Larvae often burrow in the river bottom. |
| *Stylurus laurae* | Laura's clubtail | EN | NL | Need shallow, sandy, or sandy-muddy bottomed creeks with forested shorelines. Adults use riffle areas in the stream for foraging. |
| Mammals |  |  |  |  |
| *Myotis leibii* | Eastern small-footed myotis | EN | NL | Foraging resources include wetlands and waterbodies. |
| *Myotis lucifugus* | Little brown myotis | EN | EN | Wetlands and areas around waterbodies (e.g., riparian areas and forest edges) are important foraging habitats. |
| *Myotis septentrionalis* | Northern myotis | EN | EN | Wetlands and areas around waterbodies (e.g., riparian areas and forest edges) are important foraging habitats. |
| *Perimyotis subflavus* | Tri-coloured bat | EN | EN | Wetlands and areas around waterbodies (e.g., riparian areas and forest edges) are important foraging habitats. |
| Molluscs |  |  |  |  |
| *Cyclonaias tuberculata* | Purple wartyback | TN | NL | Can be found in small to large rivers with various types of substrates. |
| *Epioblasma rangiana* | Northern riffleshell | EN | EN | Found in riffle areas within rivers or streams with rocky, sand, or gravel bottoms. |
| *Epioblasma triquetra* | Snuffbox | EN | EN | Found in rivers in shallow riffle areas. |
| *Lampsilis fasciola* | Wavy-rayed lampmussel | TN | SC | Found in riffle areas in medium to large-sized streams. |

| Species Name | Common Name | SARO | SARA | Habitat Description and Use |
| --- | --- | --- | --- | --- |
| *Ligumia nasuta* | Easter pondmussel | SC | SC | Occurs in sheltered areas of lakes, in slack-water areas of rivers and coastal wetlands. |
| *Mesodon zaletus* | Toothed globe | EN | EN | Habitat includes river bluffs and ravines, especially on steep slopes along rivers. |
| *Obliquaria reflexa* | Threehorn wartyback | TN | TN | Appears in large rivers with moderate current and in shallow embayments and reservoirs with little current. |
| *Obovaria subrotunda* | Round hickorynut | EN | EN | Found in rivers with clay, sand, or gravel bottoms and shallow areas of lakes with firm sand. |
| *Philomycus carolinianus* | Carolina mantleslug | TN | TN | Found in riparian areas of older-growth forests. |
| *Pleurobema sintoxia* | Round pigtoe | EN | EN | Found in rivers of various sizes with deep water and sandy, rocky, or muddy bottoms. |
| *Ptychobranhus fasciolaris* | Kidneyshell | EN | EN | Found in small to medium-sized rivers. It prefers shallow, clear, swift-moving water. |
| *Quadrula quadrula* | Mapleleaf | SC | SC | Found in large rivers with slow to moderate currents and firmly packed sand, gravel, or clay and mud bottoms. It also lives in lakes and reservoirs. |
| *Simpsonaias ambigua* | Salamander mussel | EN | EN | Occurs in rivers and streams with soft bottoms and swift currents. |
| *Toxolasma parvum* | Lilliput | TN | EN | Found in rivers and wetlands having soft bottoms, such as mud, sand, and silt. |
| *Truncilla donaciformis* | Fawnsfoot | EN | EN | Inhabits medium and large rivers with moderate to slow-flowing water. It usually inhabits shallow waters (1-5 m) and has been found in lakes and reservoirs. |
| *Villosa fabalis* | Rayed bean | EN | EN | Found in small tributaries, often buried among the roots of aquatic plants. |

| Species Name | Common Name | SARO | SARA | Habitat Description and Use |
| --- | --- | --- | --- | --- |
| *Villosa iris* | Rainbow | SC | SC | Prefers small to medium-sized rivers with a moderate to strong current. Found in or near riffle areas and along the edges of vegetation in water less than one metre deep. |
| *Webbhelix multilineata* | Striped whitelip | EN | EN | Inhabits wet, lowland forest at the margins of periodically flooded areas (like marshlands or swamps), or in continuously wet areas. |
| Plants |  |  |  |  |
| *Agalinis skinneriana* | Skinner's agalinis | EN | EN | Found in river deltas and prairie wetlands. |
| *Ammannia robusta* | Scarlet ammannia | EN | EN | Found in mudflats, sand beaches, edges of wetlands and ponds. |
| *Arisaema dracontium* | Green dragon | SC | NL | Found along streams, particularly maple forests and forests dominated by Red Ash and White Elm trees. |
| *Bryoandersonia illecebra* | Spoon-leaved moss | TN | EN | Swamps, marshes, and wet meadows. |
| *Buchnera americana* | Bluehearts | EN | EN | Wet meadow communities between sand dunes along shorelines. |
| *Carex lupuliformis* | False hop sedge | EN | EN | Riverine swamps and marshes, and around temporary forest ponds. |
| *Cyperus subsquarrosus* | Small-flowered Lipocarpha | TN | EN | Grows on sandy beaches that are seasonally flooded and are protected from high waves or strong currents. |
| *Cypripedium candidum* | Small white lay's-slipper | EN | TN | Grows in moist prairies, savannahs, and rich calcareous (limestone) wetlands, known as fens. |
| *Eleocharis equisetoides* | Horetail spike-rush | EN | EN | Grows in shallow water along the edges of ponds. |

| Species Name | Common Name | SARO | SARA | Habitat Description and Use |
| --- | --- | --- | --- | --- |
| *Eleocharis geniculata* | Bent spike-rush | EN | EN | Found on wet, sandy to muddy soil in open flats along the shore of Lake Erie. Occurs occasionally along the edges of wet meadows and seasonal ponds further inland. |
| *Fraxinus nigra* | Black ash | EN | NL | A wetland species found in swamps, floodplains and fens. |
| *Fraxinus quadrangulata* | Blue ash | TN | EN | Grows in deciduous floodplain forests, and along sandy beaches. |
| *Gymnocladus dioicus* | Kentucky coffee-tree | TN | TN | Often found in floodplains, woodland edges of marshes. |
| *Hibiscus moscheutos* | Swamp rose-mallow | SC | SC | Restricted to shoreline marshes. Commonly found in deep-water cattail marshes and in meadow marshes, wetlands. |
| *Justicia americana* | American water-willow | TN | TN | Occurs in fresh water, fairly open habitats with little competition from other aquatic plants. Grows along the shores and in the shallow waters of streams, rivers, lakes, ditches, and occasionally wetlands. |
| *Morus rubra* | Red mulberry | EN | EN | Occurs in floodplains and river valleys and on swales of sandspits. |
| *Phegopteris hexagonoptera* | Broad beech fern | SC | NL | Occurs in moist areas like lower valley slopes, bottomlands and swamps, stream floodplains, riparian areas. |
| *Plantago cordata* | Heart-leaved plantain | EN | EN | A semi-aquatic plant, it is found in relatively undisturbed wet woods, often along the rocky or gravelly limestone beds of shallow, slow-moving, clear streams. |
| *Platanthera leucophaea* | Eastern prairie fringed orchid | EN | EN | Grows in wetlands, fens, swamps, and tallgrass prairie. |
| *Potamogeton hillii* | Hill's pondweed | SC | SC | Found in slow-moving streams, ditches, ponds, lakes, and wetlands. It grows in clear, cold alkaline waters. |
| *Ptelea trifoliata* | Common hoptree | SC | SC | Some individuals found on beach sand. |

| Species Name | Common Name | SARO | SARA | Habitat Description and Use |
| --- | --- | --- | --- | --- |
| *Quercus shumardii* | Shumard oak | SC | NL | Can grow close to water and in swampy areas. |
| *Ripariosida hermaphrodita* | Virginia mallow | EN | EN | Found in riparian habitats that are flooded in most years. Loose sandy or rocky soils of scoured riversides and floodplains, and disturbed areas along roadsides and railroad banks are its preferred habitats. |
| *Stylophorum diphyllum* | Wood-poppy | EN | EN | Found in rich mixed deciduous woodlands, forested ravines and slopes, and along wooded streams. |
| *Symphyotrichum prenanthoides* | Crooked-stem aster | SC | SC | Grows in wet areas along the banks of rivers and streams. |
| *Tephrosia virginiana* | Virginia goat's-rue | EN | EN | Grows in open, sunny areas with sandy soil, such as sand dunes. |
| *Trillium flexipes* | Drooping trillium | EN | EN | Populations are associated with watercourses, usually on better-drained microsites on floodplain terraces or adjacent river and stream slopes. |
| *Valeriana edulis ssp. ciliata* | Hairy valerian | TN | EN | Typically found on wet and moderately wet prairies and fens. |
| Reptiles |  |  |  |  |
| *Apalone spinifera* | Spiny softshell turtle | EN | EN | Found in coastal areas and major rivers/tributaries to Lake Erie, Lake St Clair, and Lake Huron. |
| *Chelydra serpentina* | Snapping turtle | SC | SC | Spend most of their lives in water, prefer shallow waters. Suitable nesting sites are usually gravelly or sandy areas along streams. |
| *Chrysemys picta marginata* | Midland painted turtle | NL | SC | Occupy slow-moving, relatively shallow, and well-vegetated wetlands and water bodies with abundant basking sites and organic substrate. Found in association with submergent aquatic plants, which are used for cover and feeding. |

| Species Name | Common Name | SARO | SARA | Habitat Description and Use |
| --- | --- | --- | --- | --- |
| *Clemmys guttata* | Spotted turtle | EN | EN | Prefers ponds, marshes, bog, and even ditches with slow-moving, unpolluted water and an abundant supply of aquatic vegetation. Usually hibernates in wetlands. |
| *Emydoidea blandingii* | Blanding's turtle | EN | EN | Occurs in a variety of wetland habitats, slow-flowing rivers and creeks, pools, lakes, bays, sloughs, marshy meadows, and artificial channels. Will select all wetland types over lotic environments and have also shown a preference for ponds and marshes when available. |
| *Graptemys geographica* | Northern map turtle | SC | SC | Inhabits rivers and lakeshores in summer. In winter, the turtles hibernate on the bottom of deep, slow-moving sections of the river. |
| *Heterodon platirhinos* | Eastern hog-nosed snake | NL | TN | Use beach and dune habitat, often relying on driftwood and other artificial and natural ground cover. Prefer forested areas and wetlands. |
| *Pantherophis gloydi* | Eastern foxsnake | TN | NL | Typically associated with unforested habitats, including old fields, prairies, savannas, shorelines, rock barrens, marshes, and beach dunes. They exhibit a strong preference for shoreline edge habitats. |
| *Plestiodon fasciatus* | Five-lined skink | EN | NL | Occur along or near the shores of Lakes Erie, St. Clair, and Huron, and major tributaries. Individuals are generally found on stabilized sand dunes, savannahs, relic prairie patches, open forested areas, and wetland areas. Stabilized dune and savannah are strongly preferred. |
| *Regina septemvittata* | Queensnake | EN | EN | An aquatic species that is seldom found more than a few metres from the water. It prefers rivers, streams, and lakes with clear water, rocky or gravel bottoms, lots of places to hide, and an abundance of crayfish. |

| Species Name | Common Name | SARO | SARA | Habitat Description and Use |
| --- | --- | --- | --- | --- |
| *Sistrurus catenatus* | Massasauga rattlesnake | EN | EN | Lives in a variety of habitats, including tall grass prairie, bogs, marshes, shorelines, forests, and alvars. Non-pregnant females and males forage and mate in lowland habitats such as grasslands, wetlands, bogs, and the shorelines of lakes and rivers. |
| *Sternotherus odoratus* | Eastern musk turtle | SC | SC | Found in ponds, lakes, marshes, and rivers that are generally slow-moving have abundant emergent vegetation and muddy bottoms that they burrow into for winter hibernation. Nesting habitat is variable, but it must be close to the water and exposed to direct sunlight. |
| *Thamnophis butleri* | Butler's gartersnake | EN | EN | Prefers open, moist habitats, such as dense grasslands and old fields, with small wetlands where it can feed on leeches and earthworms. |
| *Thamnophis sauritus* | Eastern ribbonsnake | SC | TN | Usually found close to water, especially in marshes, where it hunts for frogs and small fish. A good swimmer, it will dive in shallow water, especially if it is fleeing from a potential predator. |

For those species whose status reports or recovery strategies provided specific wetland use information, marshes and swamps were used by more species than bogs and fens (Figure 4). River and stream habitat was predominantly used by molluscs, while lakes, ponds, and pools were predominantly used by amphibians and reptiles (Table 1). Use of habitat types by all other taxonomic groups was fairly even, with wetland habitats used for foraging by most bird and mammal species (Table 1).

### Invasive Plant Occurrence & Impact Mechanisms

A total of 33 invasive plant species were identified as current or imminent threats to aquatic and wetland habitats in the Carolinian Zone (Table 2). Twenty-six species had confirmed observations within the Carolinian based on EDDMaps data from 2014–2024 (current threats). The most frequently observed species were garlic mustard (*Alliaria petiolata*) with 4,202 observations, Japanese knotweed (*Reynoutria japonica*) with 1,045 observations, and purple loosestrife (*Lythrum salicaria*) with 900 observations (Table 2). These species are known for their aggressive spread and high environmental impact (Blossey et al., 2001; Gerber et al., 2008; Rodgers et al. 2022; Table 2). However, it should be noted that the high environmental impact associated with *L. salicaria* invasion (GLANSIS 2025) is specifically in locations where its growth is not controlled (e.g., by biocontrol agents).

**Table 2.** Observations of invasive, non-native species identified as current or imminent invaders of the Carolinian. Species with 0 observations in the Canadian Carolinian are imminent threats, those with >1 observation are current threats. For current threats and imminent threats, environmental and socio-economic impact ratings for aquatic species are sourced from Great Lakes Aquatic Nonindigenous Species Information Systems database (GLANSIS 2025) and for species not listed (NL) or with status unknown (U) by GLANSIS (indicated with *) the Minnesota Invasive Species Advisory Council (2019). L = low impact; M = moderate impact; H = high impact.

| Species Name | Common Name | Habit | Observations | Environmental Impact | Socio-economic Impact |
| --- | --- | --- | --- | --- | --- |
| *Akebia quinata* | Chocolate vine | Terrestrial | 0 | H* | NL |
| *Alliaria petiolata* | Garlic mustard | Terrestrial | 4202 | M | L |
| *Alnus glutinosa* | Black alder | Terrestrial | 196 | M | L |
| *Artemisia absinthium* | Absinthe wormwood | Terrestrial | 0 | L* | NL |
| *Butomus umbellatus* | Flowering rush | Emergent | 263 | M | U |
| *Egeria densa* | Brazilian waterweed | Submerged | 0 | H* | NL |
| *Eichhornia crassipes* | Common water hyacinth | Free-floating | 13 | H* | NL |
| *Epilobium hirsutum* | Great hairy willowherb | Emergent | 149 | M | L |
| *Frangula alnus* | Glossy buckthorn | Terrestrial | 267 | H | L |
| *Glyceria maxima* | Rough manna grass | Emergent | 23 | M | L |
| *Hydrilla verticillata* | Hydrilla | Submerged | 1 | H | H |
| *Hydrocharis morsus-ranae* | European frog-bit | Free-floating | 320 | M | M |
| *Iris pseudacorus* | Yellow flag iris | Emergent | 402 | H | H |
| *Juncus inflexus* | European meadow rush | Emergent | 0 | M | L |
| *Lysimachia vulgaris* | Yellow loosestrife | Terrestrial | 44 | M | L |
| *Lythrum salicaria* | Purple loosestrife | Emergent | 900 | H | U |
| *Marsilea quadrifolia* | European water-clover | Emergent | 1 | U; M* | L |
| *Myriophyllum spicatum* | Eurasian watermilfoil | Submerged | 108 | H | H |
| *Najas minor* | Brittle waternymph | Submerged | 7 | M | M |
| *Nasturtium officinale* | Watercress | Emergent | 48 | U; L* | L |
| *Nitellopsis obtusa* | Starry stonewort | Submerged | 0 | H | H |
| *Nymphoides peltata* | Yellow floating heart | Rooted-floating | 2 | M | M |
| *Phalaris arundinacea* | Reed canary grass | Terrestrial | 356 | H | M |
| *Phragmites australis* ssp. *australis* | European common reed | Emergent | 513 | H | M |
| *Pistia stratiotes* | Water lettuce | Free-floating | 9 | H | H |
| *Potamogeton crispus* | Curly leaf pondweed | Submerged | 174 | M | M |
| *Reynoutria japonica* | Japanese knotweed | Terrestrial | 1045 | H** | NL |
| *Robinia pseudoacacia* | Black locust | Terrestrial | 635 | H* | NL |
| *Rorippa sylvestris* | Yellow fieldcress | Terrestrial | 1 | U; L* | L |
| *Salix fragilis* | Crack willow | Terrestrial | 0 | U; L* | M |
| *Trapa natans* | Water chestnut | Rooted-floating | 5 | H | H |
| *Typha angustifolia* | Narrow-leaved cattail | Emergent | 172 | H | L |
| *Typha* × *glauca* | Hybrid cattail | Emergent | 213 | H* | NL |

\*\* Lavoie, C. (2017) The impact of invasive knotweed species (*Reynoutria* spp.) on the environment: review and research perspectives. *Biological Invasions*, 19, 2319-2337.

Mechanisms of impact (Table 3) varied to some degree by growth form, with submersed aquatic and floating invasive plant species tending to exert a greater diversity of mechanisms than terrestrial and emergent invasive plants (Figure 5). Plant species with more diverse mechanisms of impact also tended to be rated as having higher overall environmental impact (Figure 5). Although every growth form of invasive wetland plant we evaluated (terrestrial, emergent, submersed, free-floating or rooted-floating) had examples of species with high environmental impact, only terrestrial and emergent growth forms have examples rated as having low environmental impact (Figure 5).

**Figure 5.**
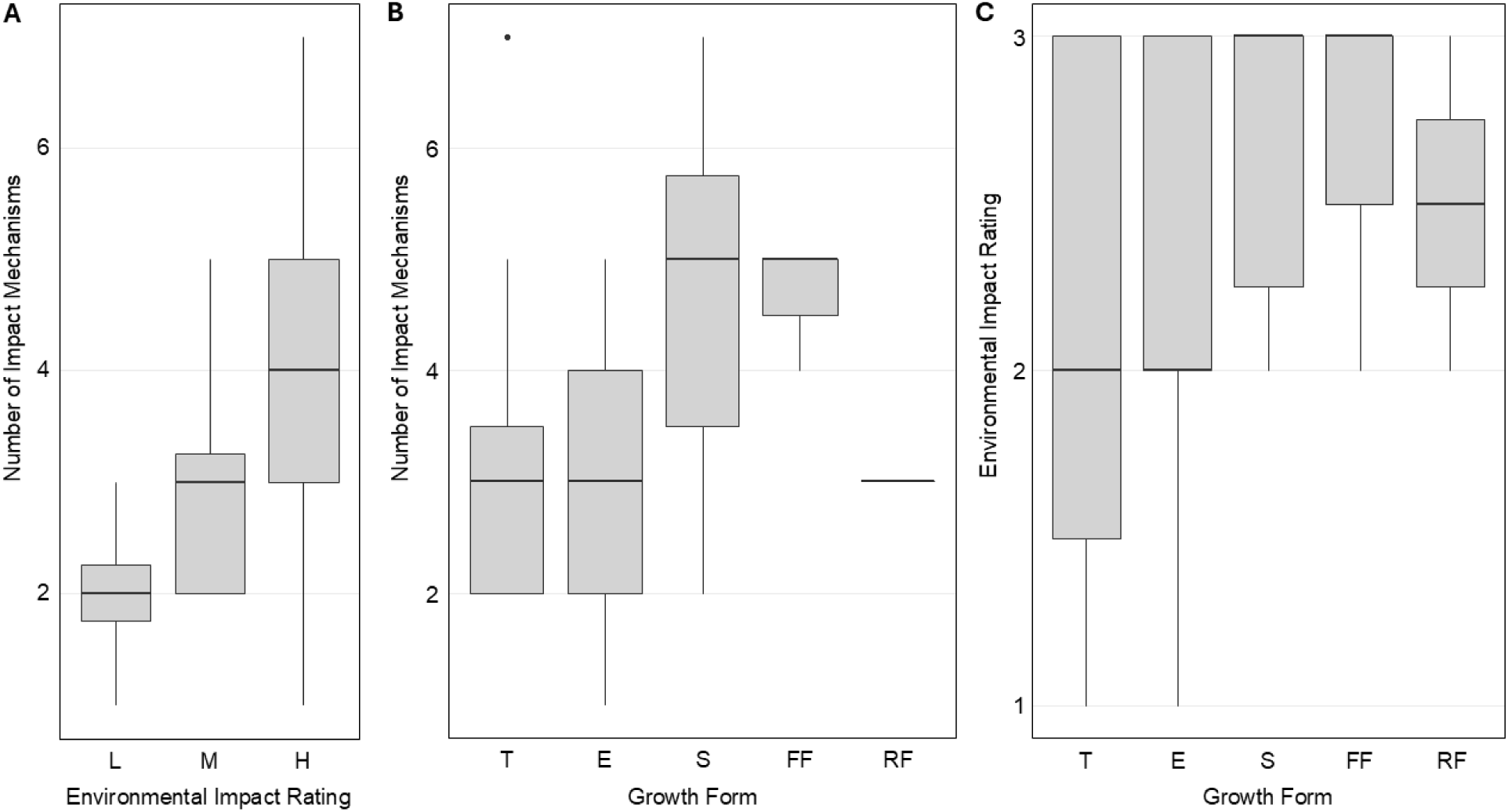
Relationships among environmental impact rating, growth form, and number of impact mechanisms for invasive species. (A) Number of impact mechanisms across GLANSIS environmental impact ratings (H = high, M = medium; L = low). (B) Number of impact mechanisms across growth form (T = terrestrial; E = emergent; S = Submersed; FF= Free-floating; RF = rooted-floating). (C) Environmental impact rating (ordinal ranking 3 = High, 2 = Medium, 1 = Low) across growth forms. Boxes show interquartile range (IQR), horizontal lines indicate medians, whiskers represent 1.5 × IQR, and points indicate outliers.

**Table 3.** Mechanisms of impact for invasive plants found in (current threats), and those that might soon invade (imminent threats) the Carolinian.

| Species Name | Common Name | Resource monopolization - Above ground | Resource monopolization - Below ground | Rapid Growth | Impenetrable stands or mats | Altered hydrology | Allelopathy | Smothering/Strangling | De-oxygenation of water | Hybridization | Altered benthic conditions | Interactions with other invasive species | Pest and disease transmission | Modification of nutrient regime | High reproductive potential | Increased erosion |
| --- | --- | --- | --- | --- | --- | --- | --- | --- | --- | --- | --- | --- | --- | --- | --- | --- |
| *Akebia quinata* | Chocolate vine | x | x | x | x |  |  | x |  |  |  |  |  |  |  |  |
| *Alliaria petiolata* | Garlic mustard |  |  |  |  |  | x | x |  |  |  |  | x |  |  |  |
| *Alnus glutinosa* | Black alder | x |  |  |  | x |  |  | x |  |  |  |  |  |  |  |
| *Artemisia absinthium* | Absinthe wormwood | x | x |  |  |  |  |  |  |  |  |  |  |  |  |  |
| *Butomus umbellatus* | Flowering rush | x | x |  |  | x |  |  |  |  |  |  |  |  |  |  |
| *Egeria densa* | Brazilian waterweed | x |  |  | x | x |  |  | x |  | x |  |  |  |  |  |
| *Eichhornia crassipes* | Water hyacinth | x | x |  | x |  |  |  |  |  | x |  |  |  |  |  |
| *Epilobium hirsutum* | Great hairy willow herb | x |  | x |  |  |  |  |  |  |  |  |  |  |  |  |
| *Frangula alnus* | Glossy buckthorn | x |  |  |  |  |  |  |  |  |  |  | x |  | x |  |
| *Glyceria maxima* | Rough manna grass | x |  |  | x | x |  |  | x |  |  |  |  |  |  |  |
| *Hydrilla verticillata* | Hydrilla | x | x | x | x | x |  |  | x |  | x |  |  |  |  |  |
| *Hydrocharis morsus-ranae* | European frog-bit | x | x | x | x |  |  |  | x |  |  |  |  |  |  |  |
| *Iris pseudacorus* | Yellow flag iris |  |  |  |  |  | x |  |  |  |  |  |  |  |  |  |
| *Juncus inflexus* | European meadow rush | x | x |  |  |  | x |  |  |  |  |  |  |  |  |  |
| *Lysimachia vulgaris* | Yellow loosestrife |  | x |  |  | x |  |  |  |  |  |  |  |  |  |  |
| *Lythrum salicaria* | Purple loosestrife |  | x | x |  |  |  |  |  |  |  |  |  | x |  |  |
| *Marsilea quadrifolia* | European water-clover | x | x |  |  |  |  |  |  |  |  |  |  |  |  |  |
| *Myriophyllum spicatum* | Eurasian watermilfoil | x |  | x | x |  |  | x |  | x |  | x |  |  |  |  |
| *Najas minor* | Brittle waternymph |  |  |  |  |  |  |  | x |  |  | x |  |  |  |  |
| *Nasturtium officinale* | Watercress |  |  |  |  | x |  |  |  |  |  |  |  |  |  |  |
| *Nitellopsis obtusa* | Starry stonewort |  |  |  |  |  | x | x |  |  | x |  |  |  |  |  |
| *Nymphoides peltata* | Yellow floating heart | x | x | x |  |  |  |  |  |  |  |  |  |  |  |  |
| *Phalaris arundinacea* | Reed canary grass | x | x | x | x |  |  |  |  |  |  |  |  |  |  |  |
| *Phragmites australis* | European common reed | x | x |  | x | x |  |  |  |  |  |  |  |  |  |  |
| *Pistia stratiotes* | Water lettuce | x |  | x |  |  |  | x |  |  |  | x | x |  |  |  |
| *Potamogeton crispus* | Curlyleaf pondweed | x | x | x |  |  | x |  |  | x |  |  |  |  |  |  |
| *Reynoutria japonica* | Japanese knotweed | x | x | x | x |  | x |  |  |  |  |  |  |  | x | x |
| *Robinia pseudoacacia* | Black locust |  | x |  |  |  |  |  |  |  |  |  |  | x |  |  |
| *Rorippa sylvestris* | Yellow fieldcress |  |  |  |  |  | x |  |  | x |  |  |  |  |  |  |
| *Salix fragilis* | Crack willow | x |  |  |  | x |  |  |  |  |  |  |  |  |  | x |
| *Trapa natans* | Water chestnut | x | x |  |  |  |  | x |  |  |  |  |  |  |  |  |
| *Typha angustifolia* | Narrow-leaved cattail | x | x | x |  |  |  |  |  | x |  |  |  |  |  |  |
| *Typha x glauca* | Hybrid cattail | x | x | x | x |  |  |  |  | x |  |  |  |  |  |  |
| Total |  | 24 | 19 | 13 | 11 | 9 | 7 | 6 | 6 | 5 | 4 | 3 | 3 | 2 | 2 | 2 |

Aboveground (24 of 33 invasive wetland plant species) and belowground resource monopolization (19 of 33) are the most common impact mechanisms among the 33 invasive wetland plants we evaluated, inferring that the threat invasive wetland plants pose to species at risk plants is principally via indirect competition for resources. However, the related mechanisms of rapid growth (13 of 33) resulting in smothering/strangling (6 of 33) and the creation of impenetrable stands or mats (11 of 33) are also very common mechanisms among the invasive wetland plants presenting a current or imminent threat to the Carolinian, suggesting pre-emptive competition for space may also be important.

Mechanisms like alleopathy (7 of 33) and hybridization risk (5 of 33) are comparatively less common means of effecting native plant species, but the species of plants exhibiting these mechanisms are among the most commonly observed in the Carolinian. For example, *A. petiolata* (4202 occurrences), *R. japonica* (1045 occurrences), and Yellow iris (*Iris pseudacorus*; 402 occurrences) are all capable of allelopathy, while *T.* x *glauca* (213 occurrences), Curlyleaf pondweed (*Potamogeton crispus*; 174 occurrences) Narrow-leaved cattail (*T. angustifolia*; 172 occurrences), and *M. spicatum* (108 occurrences) all present a threat to native members of their genera through hybridization (Table 3). Because the threat of hybridization is restricted to species that are closely related to the invader, we recognize this impact mechanism is of relatively minor threat to species at risk plants in the Carolinian. However, we note that this may be an important mechanism behind broader ecological effects of plant invasion. The principal impact mechanism by which invasive wetland plants in our analysis might threaten at-risk plants is indirect resource competition or perhaps pre-emptive competition for space.

The mechanisms of impact by which invasive wetland plants threaten animal species at risk are almost entirely related to habitat modification through different means (Table 3). Most common is by creating dense, impenetrable vegetation mats or stands (11 of 33 species), although several invasive plants are documented to alter hydrology (9 of 33), benthic conditions (4 of 33), or nutrient regimes (2 of 33). Six of 33 species also contributed to de-oxygenation of water, mainly submersed and floating species such as *H. morsus-ranae*, Brazilian waterweed (*Egeria densa*), and Hydrilla (*Hydrilla verticillata*). Thus, the principal mechanisms by which invasive wetland plants impact animal species at risk is via degradation of habitat quality.

Of the seven species not yet observed in the Carolinian (imminent threats), species with terrestrial growth forms were most common (57%). Of those species found in the Carolinian (current threats), emergent species (42%) and terrestrial species (31%) were the most common observed growth forms. Both *P. australis* and narrow-leaved cattail (*Typha angustifolia)* were among the most widespread emergent species. Both have high environmental impacts caused by forming dense monocultures that displace native vegetation, altering hydrology and modifying wetland structure and function (Bansal et al. 2019; Hazelton et al., 2014). Hybrid cattail (*Typha* x *glauca*) is a hybrid of native T*ypha latifolia* and invasive *T. angustifolia* that likely has similarly high environmental impacts: it grows in dense monocultures that increase methane emissions (Lawrence et al., 2016a), and negatively impact macroinvertebrate abundances (Lawrence et al., 2016b) and fish diversity (Schrank and Lishawa, 2019).

Several submerged and free-floating species, such as Watercress (*Nasturtium spicatum*) and European frog-bit (*Hydrocharis morsus-ranae*), were also present, but in lower numbers. It warrants noting that because these occurrence data come from EDDMapS, they may reflect lower survey effort in open water habitats where floating and submersed growth forms occur. Although community science platforms such as iNaturalist and EDDMaps can provide valuable data for early detection and monitoring of invasive plants, these platforms are inherently biased toward accessible and frequently visited areas (e.g., Guerts et al., 2023). This bias may be an advantage in invasive plant surveillance, as invasive species often distribute preferentially in human-affected and accessible areas like roadways, pastures, or developed areas (Lázaro-Lobo et al. 2021); however, invasive plants with submersed aquatic growth forms may escape notice relative to more visible emergent and floating plant species. Anecdotally, the first confirmed observation of *Hydrilla verticillata* in Canada in July 2024 in the Hillman Marsh Conservation Area (McQuaid, 2024) was posted to iNaturalist within weeks of its independent discovery by academic researchers (Rooney pers. comm.). Davidson et al. (2021) emphasize the importance of community monitoring approaches (aka participatory or citizen science), particularly for aquatic plants, which are underrepresented in many current monitoring programs in the Great Lakes region. Larson et al. (2020) noted the value of community monitoring programs like iNaturalist in the early detection of invasive species. However, for inaccessible areas with high connectivity to invaded habitats there is a need for formal, targeted monitoring protocols that prioritize early detection and adequately resourced long-term monitoring to accurately evaluate management efforts.

### Distribution Overlap of Species At Risk and Invasive Plants

The most striking outcome of our analysis is the high spatial overlap between the occurrence of invasive wetland plants and the occurrence (Figure 6) and richness (Figure 7) of species at risk in the Carolinian zone. This overlap is spatially aggregated in the coastal wetlands of Point Pelee National Park, Rondeau Provincial Park, and the Long Point World Biosphere Reserve along the north shore of Lake Erie, where a high occurrence and richness of species at risk is documented. In these conservation lands, our results highlight a high occurrence of invasive plants, including those ranked as having moderate environmental impact, such as European frog-bit (*Hydrocharis morsus-ranae*) and Flowering rush (*Butomus umbellatus*), as well as those ranked as having high environmental impact, such as European common reed (*Phragmites australis*), Eurasian watermilfoil (*Myriophyllum spicatum*), and hybrid cattail (*Typha* x *glauca;* Figure 8). In recognition of the threat invasive plants posed to species at risk in these three protected coastal wetlands, all three of these locations have undertaken invasive *P. australis* control programs in recent years. Rondeau and Long Point participated in an Emergency Use Registration pilot project to use herbicide to control *P. australis* in standing water (Robichaud and Rooney, 2023) and Point Pelee undertook costly but effective manual removal (Ward et al., 2025).

**Figure 6.**
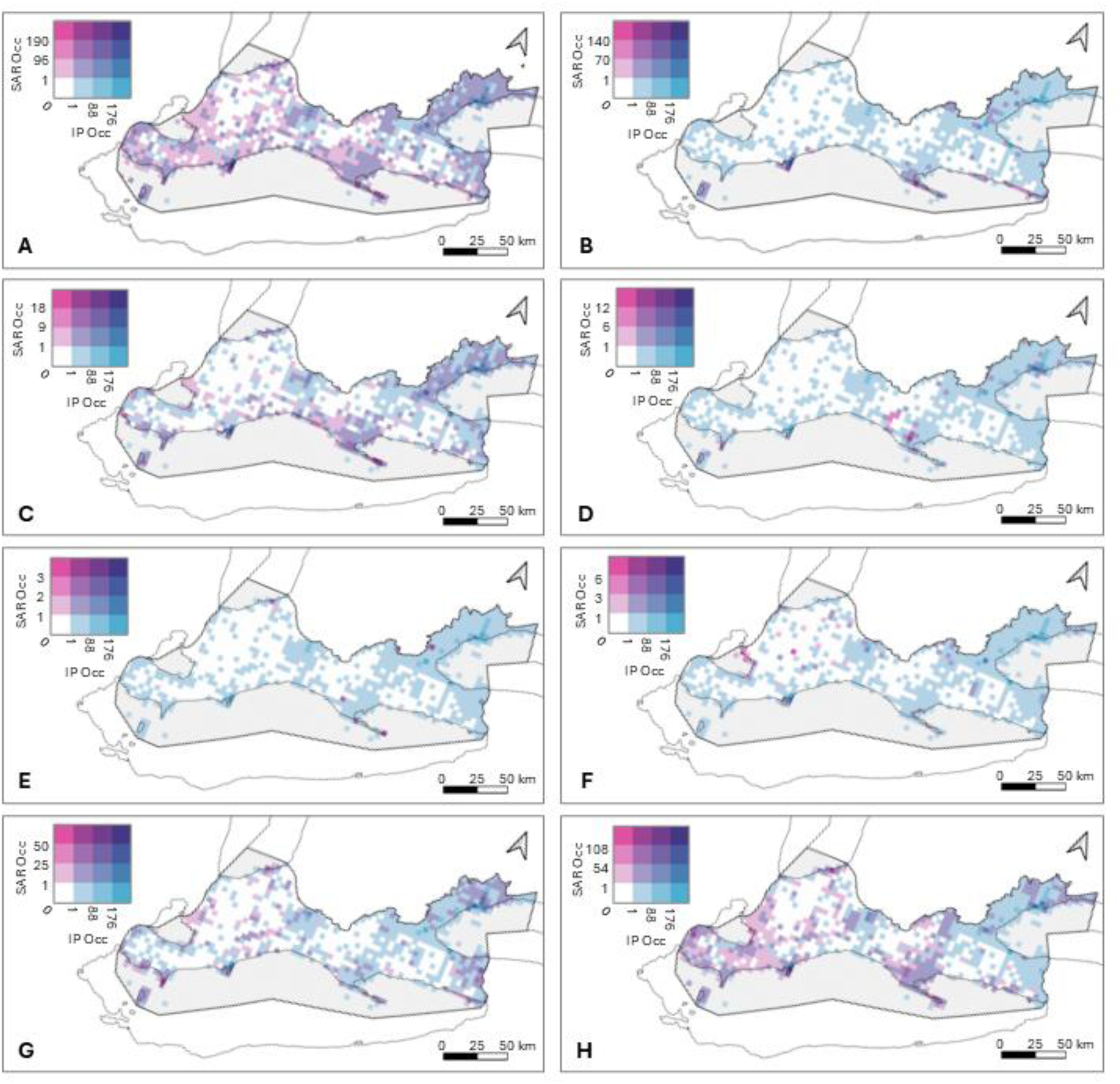
Choropleth of invasive wetland plant species occurrence overlaid with at-risk species occurrence for: (A) all at risk species groups, (B) amphibians, (C) birds, (D) insects, (E) mammals, (F) mollusks, (G) plants, (H) reptiles.

**Figure 7.**
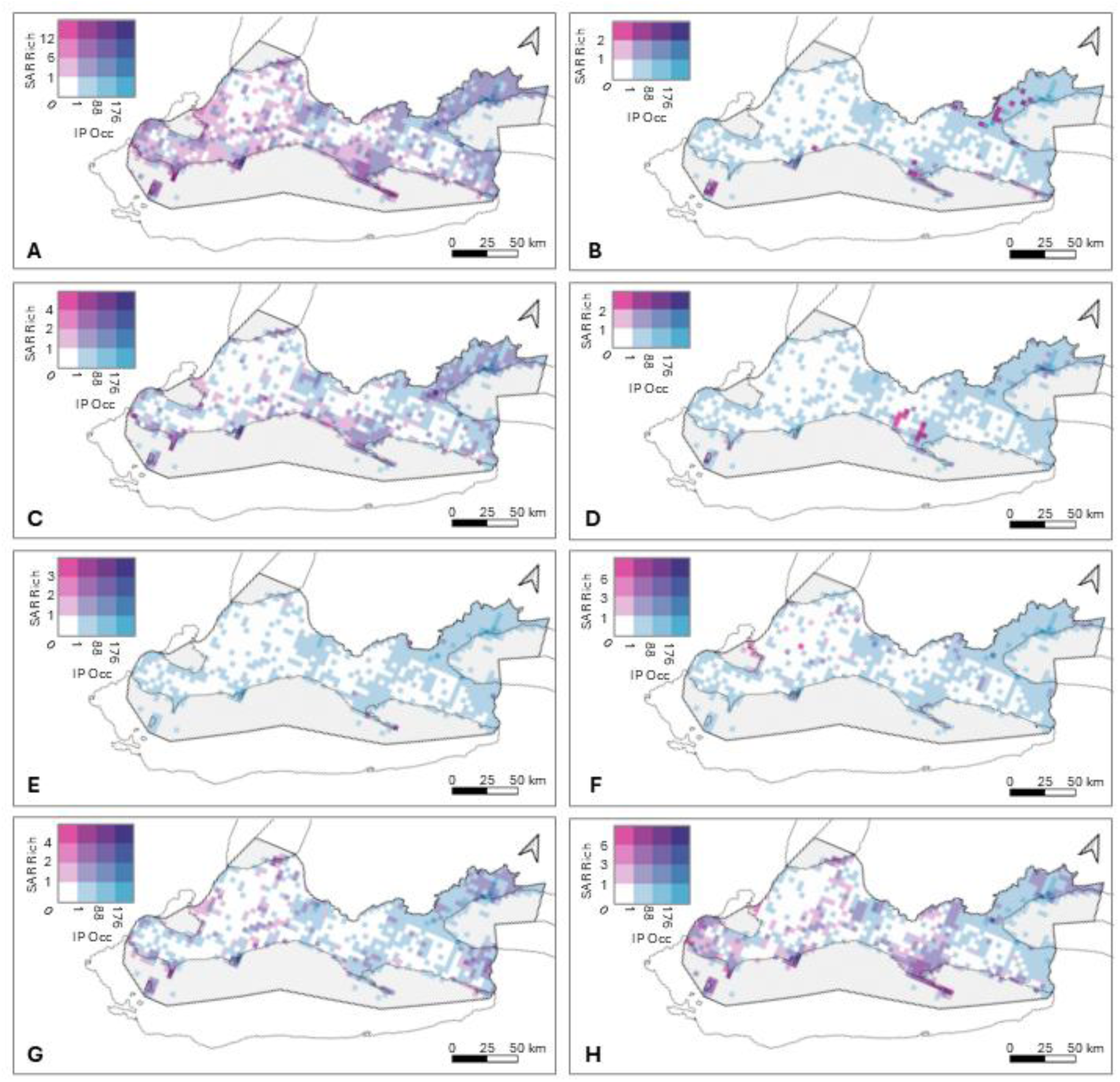
Choropleth of invasive wetland plant species occurrence overlaid with at-risk species richness for: (A) all at risk species groups, (B) amphibians, (C) birds, (D) insects, (E) mammals, (F) mollusks, (G) plants, (H) reptiles.

**Figure 8.**
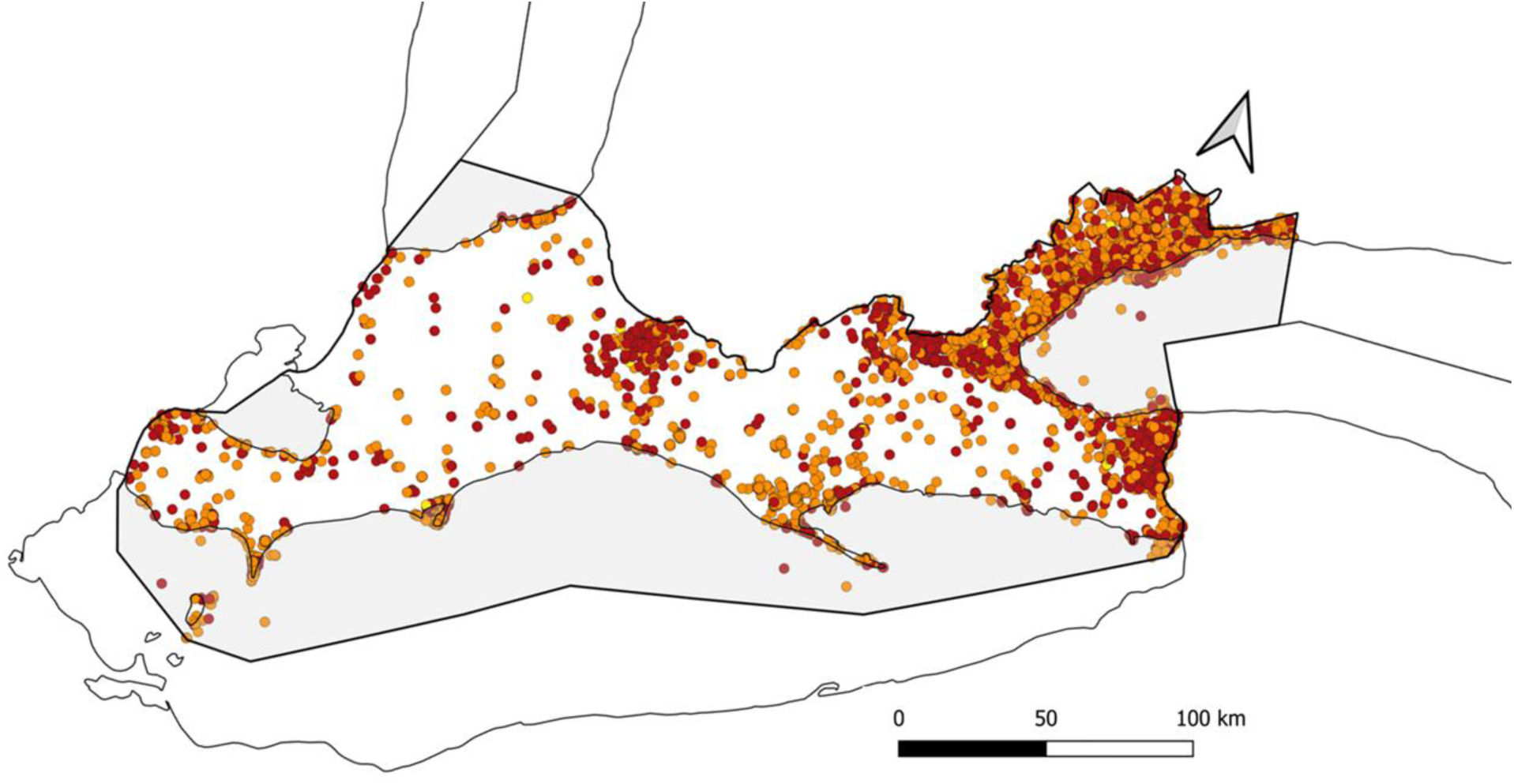
Occurrence of identified invasive plant species with environmental impact indicated by colour (red = High environmental impact, orange = Moderate environmental impact, yellow = Low environmental impact). Sources of environmental impact classes are following Table 2.

Yet our analysis identifies other hotspots of risk and vulnerability that have not previously been highlighted as conservation priorities. These other hotspots of co-occurrence between invasive wetland plants and species at risk occurrence and richness are found in urban centres with higher human population densities and activity. For example, across the Greater Toronto Area surrounding northwestern Lake Ontario, along the Niagara-Buffalo corridor connecting Lake Ontario and Lake Erie, near the Detroit-Windsor border south of Lake St. Clair, from Sarnia to Grand Bend along the southern shore of Lake Huron, and around the City of London (Figure 1). The greater occurrence of invasive wetland plants in areas of higher human population density is perhaps unsurprising, given the well-documented relationship between invasive aquatic plants and human land use (e.g., Zedler and Kercher, 2004; Trebitz and Taylor, 2007). What is more surprising is that, despite lacking the protections afforded Long Point, Rondeau and Point Pelee, these areas also support relatively high richness and occurrence of species at risk, especially of birds, (Figures 6c, 7c), plants (Figures 6g, 7g), and reptiles (Figures 6h, 7h). We recommend that these areas of high co-occurrence of species at risk and invasive wetland plants be the subject of strategic conservation planning in Ontario. Habitat protection for the conservation of at-risk species evidently must include wetland and semi-aquatic habitats in urban areas of the Carolinian, and invasive plant management in these urban habitats may prove essential for their protection.

Another area emerging as a monitoring priority is the wetland complex northeast of Lake St. Clair: the land traditionally called Bkejwanong, an Anishinaabemowin name meaning “where the waters divide,” which is stewarded by the Walpole Island First Nation (Fehr, 2011). This area supports a high occurrence and diversity of at-risk amphibians, birds, plants, reptiles and molluscs (Figures 6f, 7f). Yet unlike other large, protected wetlands in the Carolinian (Point Pelee, Rondeau, and Long Point), Bkejwanong has relatively low invasive plant occurrences reported in EDDSmaps. This could be under-reporting due to restricted access or low engagement with EDDSmaps, but regardless, we highlight this territory as a high priority for invasive species monitoring – either to address data gaps or to ensure continued protection from invasion through early detection.

Species at risk that exclusively or primarily use coastal wetland and shallow open water habitats most overlapped with aggressive, monoculture-forming invasives, including *P. australis* and *T.* x *glauca*. The hydrological connectivity of coastal wetlands facilitates the movement of invasive plant propagules and puts these habitats at greater risk of invasion than inland systems. Our analysis is consistent with regional assessments, including the State of the Great Lakes (Environment and Climate Change Canada, 2022), which has cited invasive plants as some of the most ecologically impactful invasive species in the Great Lakes. The GLANSIS database identifies *P. australis* as one of the most widespread and impactful invaders, with its range continuing to expand northward along transportation corridors and disturbed shorelines (US Environmental Protection Agency, 2023).

These invasion patterns are concerning for species at risk that depend on structurally diverse and heterogeneous wetland habitats. For example, *P. australis-*invaded habitat excludes many marsh-nesting birds (Robichaud and Rooney, 2017), like the Least bittern, Virginia rail, and King rail, which require a mosaic of interspersed emergent vegetation and open water for nesting and foraging (Conway and Gibbs, 2011; Lor and Malecki, 2006). Additionally, dense mats and stands of invasives (like *P. australis*) reduce the diversity and abundance of invertebrates, impacting food availability for many wetland birds that rely on aquatic and emergent invertebrates during breeding and migration (Gratton and Denno, 2005; Talley and Levin, 2001). Aquatic reptiles require basking sites for thermoregulation; tall, fast-growing invasives could degrade basking habitat as they block sunlight at ground level (Stratoulias and Tóth, 2020). The dense mats and stands of invasives can also hinder reptile movement as well as limit access to and availability of nesting sites (Vlk et al. 2021). Further, changes to soil moisture and temperature caused by shading, root water uptake, and changes in evapotranspiration (He, 2014) can negatively impact suitability of nesting sites and egg incubation (e.g., Schwarzkopf and Brooks, 1987; Bolton and Brooks 2010). Indeed, our spatial analysis reveals that wetland birds (Short-eared owl, Black tern, Least bittern, King rail, and Red-necked phalarope) and aquatic reptiles (Spotted turtle, Blanding’s turtle, Eastern foxsnake, Massasauga rattlesnake, Eastern musk turtle) exhibit the highest degree of overlap with invasive plant species in the Carolinian zone (Figures 6, 7), particularly in coastal marsh. They also exhibit moderate overlap in riparian habitats, where dense emergent invasives such as *P. australis* and *T.* x *glauca* may degrade breeding, nesting, and foraging habitats through shading, altered hydrology, and structural changes (He, 2014; Stratoulias and Toth, 2020; Schwarzkopf and Brooks, 1987; Vlk et al. 2021). Thus, at-risk marsh birds and reptiles may be both most exposed to invasive wetland plants and the most vulnerable to the mechanisms of impact that invasive wetland plants impose (Table 3).

At-risk aquatic plants (Spoon-leaved moss, False hop sedge, Kentucky coffee tree, Swamp rose-mallow, Horsetail spikerush and Bent spikerush) also exhibited high spatial overlap with the distribution of invasive wetland plants (Figures 6, 7) in coastal marsh. Native wetland plants are susceptible to displacement by invasive plants due to their limited dispersal capabilities and competition for limited resources (Dalle Fratte et al. 2019). Invasive monocultures like *P. australis* suppress native seed banks and reduce light availability, further disadvantaging species like Swamp rose-mallow (Farney and Bookhout, 1982; Jaworski et al., 1979). Additionally, some studies suggest that dense stands of invasives can reduce pollinator visitation to native plants, which can negatively affect reproduction (Herron-Sweet et al., 2016; Stout and Morales, 2009). Thus, species at risk aquatic plants also represent a case of high vulnerability to invasive wetland plant impacts and high exposure.

Although molluscs are one of the groups with the most at-risk species (Figure 3), they are mainly found in fast-moving, inland rivers and streams, putting them at least risk from invasive wetland plants in our study (Figures 6, 7). Invasive plants would be most impactful to mollusks if the plant species alters water flow, oxygen levels, or benthic conditions critical for mollusk survival (COSEWIC, 2006; 2008; 2013). This particular vulnerability suggests that the submersed invasive plants *E. densa* and *H. verticillata* represent serious imminent threats to at-risk mollusks, as their impact mechanisms include benthic modification, deoxygenation of the water column, and hydrologic alteration (Table 3).

Aquatic insects, particularly those associated with riverine systems, exhibit strong overlap with invasive species along major rivers like the Grand and Thames; however, insect species assessed in this study use a variety of habitats, and of the wetland habitats occupied, they are often found in inland creeks, rivers, or streams that are less invaded by wetland plants.

Mammals primarily forage in forested riparian areas where canopy cover, understory complexity, and proximity to water provide critical resources for shelter and feeding. While invasive plants are less dominant in densely forested riparian areas compared to open wetlands, they can still pose indirect threats. For example, invasives may alter understory structure, reduce native plant diversity, and disrupt trophic interactions, potentially affecting food availability and predator-prey dynamics. Although mammals are less directly impacted by invasives than other taxa assessed in this study, their reliance on riparian corridors means that degradation of these systems could have cascading effects on population viability (e.g., Lin et al., 2022).

### Management Priorities: Integrating Control and Recovery

We anticipated that mapping the overlap in distribution of invasive wetland plant occurrences and the occurrences and richness of species at risk would identify regions to prioritize in conservation and monitoring. Indeed, our analysis highlights several geographic areas within the Carolinian zone that should be prioritized for invasive plant management and species at risk recovery. Unsurprisingly, the biodiverse coastal wetlands along Lake Erie—including Long Point, Rondeau Bay, Point Pelee, and Turkey Point—exhibited the highest overlap between invasive plant abundance and species at risk occurrences, particularly for marsh-nesting birds, aquatic reptiles, and rare wetland plants (Figures 6, 7). These areas are highly connected hydrologically, facilitating the spread of invasive propagules and increasing the vulnerability of native biota. Our analysis also identified new, unexpected areas to prioritize for invasive wetland plant management where invasive wetland plants overlap with a surprising number of species at risk: the Greater Toronto Area, the Niagara-Buffalo corridor, the Detroit-Windsor border, from London to Brantford, and from Sarnia to Grand Bend. The Windsor to Point Pelee corridor, where *Hydrilla verticillata* was recently detected (McQuaid, 2024), represents an urgent priority for early detection and containment as does Bkejwanong, the lands stewarded by the Walpole Island First Nation where invasive wetlands plants seem less common. Inland areas such as the Pinery Provincial Park and the Long Point–Walsingham Forest complex also warrant attention due to their high species at risk density and documented infestations of *Phragmites australis*. Notably, several grid cells in our choropleth maps showed no observations of either SAR or invasive plants, suggesting potential data gaps rather than true absence. These undersampled areas should be targeted for formal surveys to ensure threats are not overlooked and to refine conservation priorities based on complete spatial data.

The co-existence of species at risk and invasive plants within the same habitats creates an interesting management challenge where the risk of damage from invasive plant control methods to species at risk and species at risk habitat must be considered carefully. To mitigate the risk of non-target effects from broad-spectrum approaches—like mechanical mowing or non-selective herbicide application—targeted but more labour-intensive approaches could be used in areas with lower invasive plant footprint/intensity and/or greater Species at Risk presence (Wade and Grant, 2022). However, for large, intense infestations broad-spectrum approaches may be the only feasible options for invasive plant removal and habitat recovery. For example, due to the large area infested by *P. australis* in Long Point, Lake Erie coastal marshes (over 3,500 ha), an Emergency Use Registration was issued for the first Canadian herbicide-based control program to suppress *P. australis* in standing water (Robichaud and Rooney 2021). This program resulted in significant reductions in *P. australis* stem density (>95%) (Robichaud and Rooney 2021a) with herbicide concentrations remaining below thresholds of ecological concern (Robichaud and Rooney 2021b).

Despite the threat invasive plants pose to species at risk, most federal and provincial status reports and recovery strategies do not identify specific invasive plant species of concern. This limits the ability of recovery teams to effectively implement control measures as they are often species-specific. Without species-specific monitoring data, management efforts risk being applied at the wrong time, with inappropriate methods, or in unsuitable habitat conditions. Species at risk recovery strategies must integrate invasive species control as a core component to facilitate population recovery. To support this, we recommend that recovery teams draw on existing technical guidance developed by invasive species experts. The Ontario Invasive Plant Council’s Best Management Practices series provides species-specific protocols for controlling invasive plants in sensitive habitats, like wetlands. Ultimately, strengthening inter-agency coordination between Environment and Climate Change Canada, Parks Canada, the Department of Fisheries and Oceans, the Ontario Ministry of Environment, Conservation and Parks and the Ontario Ministry of Natural Resources and Forestry will be essential to align species at risk recovery efforts with invasive species control mandates in Canada’s biodiverse Carolinian zone.

## Conclusions

Wetland and semi-aquatic habitats in the Carolinian zone of southern Ontario support over 40% of the species at risk found within the province. Marshes and shallow, open water habitats are used by many species at risk but also have the greatest abundance of high-ecological impact wetland plants, like *P. australis* and *T.* x *glauca*. This overlap is greatest for birds, reptiles, and wetland plants which are particularly threatened by habitat degradation caused by invasive plants due to their spatial overlap with invasive wetland plants and their ecological vulnerability to the mechanisms of impact used by invasive wetland plants. Mechanism of resource competition and pre-emption for space were the most common vulnerability of at-risk plants, whereas at-risk animals were vulnerable to habitat alteration, particularly through the creation of dense stands and mats, but also change to hydrology and water quality. To improve the precision of conservation actions, future recovery strategies should incorporate standardized, ecologically meaningful habitat classifications and identify specific invasive plant threats and their mechanisms of impact. This will enable more targeted and effective management interventions. Spatially explicit analyses, like the one presented here, are valuable tools for identifying priority areas for invasive species control and species at risk recovery, especially in fragmented and highly invaded landscapes like the Canadian Carolinian zone. In our case, this identified key developed areas not previously recognized as supporting species at risk under threat from invasive wetland plants. Without spatially explicit management approaches, the compounding effects of invasive plants and habitat loss will continue to weaken recovery efforts for species at risk in Canada’s most biodiverse region.

## Acknowledgements

We thank Heather Braun, Britney MacLeod, and Madeline Sutton (Canadian Wildlife Service – Environment and Climate Change Canada) for their invaluable partnership throughout this research. Their on-the-ground expertise and deep understanding of federal conservation policy were instrumental in shaping the scope and direction of this study. For data curation and organization, we thank Shafein Bedi and Adrienne Mason. For support in compiling and vetting the invasive plant list, we thank Eric Cleland (Nature Conservancy of Canada), Paul Gagnon (Long Point Region Conservation Authority), and Francine MacDonald (Ontario Ministry of Natural Resources and Forestry).

## Data Licenses

Provincially Tracked Species Data provided by Ontario Ministry of Natural Resources and Forestry, copyright: Queen’s Printer for Ontario [September 22, 2022]. Data are available by request to the Natural Heritage Information Centre. Data on invasive plant species was obtained from the open-access database EDDmapS and is available on their website (https://www.eddmaps.org/)

